# The visual ecology of a color polymorphic reef fish: the role of aggressive mimicry

**DOI:** 10.1101/2020.03.03.970988

**Authors:** Michele ER Pierotti, Anna Wandycz, Pawel Wandycz, Anja Rebelein, Vitor H Corredor, Juliana H Tashiro, Armando Castillo, William T Wcislo, W Owen McMillan, Ellis R Loew

**Affiliations:** Smithsonian Tropical Research Institute, Balboa, Panama; Department of Anatomy, Institute of Zoology, Jagiellonian University, Krakow, Poland; Faculty of Geology, Geophysics and Environment Protection, AGH University of Science and Technology, Krakow, Poland; Thunen Institute, Braunschweig, Germany; Department of Experimental Psychology, Psychology Institute, University of São Paulo, Brazil; Department of Biomedical Sciences, Cornell University, USA

**Keywords:** mimicry, communication, visual signals, sensory biology, coral reef fish, predation

## Abstract

Since all forms of mimicry are based on perceptual deception, the sensory ecology of the intended receiver is of paramount importance to test the necessary precondition for mimicry to occur, i.e. model-mimic misidentification, and to gain insight in the origin and evolutionary trajectory of the signals. Here we test the potential for aggressive mimicry by a group of coral reef fishes, the color polymorphic *Hypoplectrus* hamlets, from the point of view of their most common prey, small epibenthic gobies and mysid shrimp. We build visual models based on the visual pigments and spatial resolution of the prey, the underwater light spectrum and color reflectances of putative models and their hamlet mimics. Our results are consistent with one mimic-model relationship between the butter hamlet *H. unicolor* and its model the butterflyfish *Chaetodon capistratus* but do not support a second proposed mimic-model pair between the black hamlet *H. nigricans* and the dusky damselfish *Stegastes adustus*. We discuss our results in the context of color morphs divergence in the *Hypoplectrus* species radiation and suggest that aggressive mimicry in *H. unicolor* might have originated in the context of protective (Batesian) mimicry by the hamlet from its fish predators.

## INTRODUCTION

Despite 150 years of research since Bates’ (1862) and Wallace (1869)’s original insights, the unequivocal identification of new cases of mimicry, their evolutionary dynamics and the very definition and boundaries of the concept of mimicry are still challenging and hotly debated issues among evolutionary biologists (e.g. Vane-Wright 1980; Moynihan 1981; Ruxton et al. 2004; Rainey and Grether 2007; Wickler 2013; Dalziell and Welbergen 2016). A key realization that has emerged from the ongoing debate was that the perceptual system of the signal receiver must be put at the center of any analysis on the origins and maintenance of a mimicry system (Cuthill and Bennett 1993; Dittrich et al. 1993). Indeed, testing hypotheses of mimic-model relationships can be misleading without an appropriate eye-of-the-beholder approach (Dittrich et al. 1993). This is because the evolution of a mimic signal is shaped not by similarity with the model but by the receiver’s percepts of both the signals from model and mimic. Reducing the difference between those percepts below the receiver’s threshold for detecting a just noticeable difference (sensu Fechner 1860) will ensure perfect mimicry.

It follows that high fidelity might not be the most important requirement for efficient mimicry. Cognitive processes such as generalization (Darst and Cummings 2006; Ham et al. 2006), categorization (Chittka and Osorio 2007) and overshadowing (Mackintosh 1976), by which a conflict in the perception of multiple cues leads to only a subset of characteristics of the signal being considered at the expense of others, are likely to affect mimetic accuracy. By acting on the receiver percept of the mimic phenotype and not on the phenotype itself, selection will frequently affect only a limited subset of traits in the mimic, those most salient for the sensory system of the intended receiver. Indeed, selection might even drive the evolution of ‘imperfect’ mimics with higher mimicry performance than high fidelity mimics, for example by enhancing salient signals beyond the value characteristic of the model in a Mullerian complex, to increase effectiveness of recognition, memorization or more effective receiver manipulation (Kilner et al. 1999). In addition, a receiver percept results from alterations of the signal as it travels through the medium (e.g. air, water) from the model or mimic to the receiver’s sensory system. Characteristics of the medium (e.g. its general physical properties or those of the background against which the model/mimic are seen or heard) might enhance or attenuate certain components of the signal making perfect imitation of the model unnecessary. In conclusion, evidence in support of a particular putative mimic-model relationship needs to be grounded in an understanding of the receiver’s perceptual system and its sensory environment.

An intriguing putative case of mimicry is represented by the *Hypoplectrus* hamlet complex, a group of coral reef fish with strikingly distinct color patterns. Despite assortative mating by color morph, which led various authors to recognize them as separate species, hamlets exhibit otherwise little morphological and genetic differentiation between morphs at any one locality (Graves and Rosenblatt 1980; McCartney et al. 2003; Whiteman and Gage 2007; Puebla et al. 2008, 2014; Aguilar-Perera and González-Salas 2010). Indeed, in a recent genome-wide analysis, hamlet species only consistently differed from each other at genomic regions that contained loci implicated in the production or perception of color pattern (Hench et al. 2019). Various authors have suggested that the hamlets’ exceptional color diversity might be the result of aggressive mimicry of a number of non-predator model species by different hamlet morphs (Randall and Randall 1960; Thresher 1978; Fischer 1980; Domeier 1994; Whiteman et al. 2007; Holt et al. 2008; Puebla et al. 2018). Here, the mimic species takes the appearance of a non-predatory species in order to get close to a potential prey, small fish and epibenthic invertebrates, without eliciting an escape reaction. According to some definitions of aggressive mimicry, hamlets in fact might rather be a case of camouflage, since the mimic is signaling neither a fitness cost (as in Batesian/Mullerian mimicry) nor a benefit (as in aggressive mimicry, e.g. lures) to the receiver.

For those hamlet morphs considered mimics, one or more candidate models have been proposed, typically co-occurring herbivore, corallivore or spongivore fish species, harmless to a local prey and exhibiting various degrees of resemblance, as judged by a human viewer, to the corresponding hamlet morph (Randall and Randall 1960; Thresher 1978; Fischer 1980; Domeier 1994; Puebla et al. 2007). However, the plausibility of aggressive mimicry in hamlets rests only on these apparent color pattern similarities. A notable exception is the work of Puebla and coworkers on a butter hamlet *H. unicolor* population in Panama (Puebla et al. 2007; Puebla et al. 2018). The authors showed that the proportion of butter hamlet strikes towards their prey was significantly higher when associating with their putative model, the four-eye butterflyfish *Chaetodon capistratus*, than when striking alone, suggesting a possible fitness advantage consistent with aggressive mimicry in the butter hamlet.

Despite frequent reference to aggressive mimicry as an evolutionary engine of hamlet diversification (Thresher 1978; Fischer 1980; Domeier 1994; Puebla et al. 2007), we still lack a basic understanding of hamlet preys’ visual abilities and their potential for effective discrimination of predatory hamlet color morphs from harmless (putative) models. Here we consider three widely distributed hamlet species, the butter hamlet *(H. unicolor)*, the black hamlet *(H. nigricans)* and the non-mimic barred hamlet *(H. puella)*. We examine the visual system of two of their most common prey, namely an epibenthic coral reef fish and an open-water mysid shrimp, both in terms of color vision and visual acuity. Using spectral reflectance measurements of equivalent patches on each hamlet morph and their putative models and modeling of preys’ visual sensitivity and acuity, we gain insight into the potential for deception of each mimic hamlet morph through the eyes of their prey.

## MATERIALS AND METHODS

### Study site and species

Field work was conducted in the Bocas del Toro Archipelago, Panama, on the same reef complex (Punta Caracol; GPS 9° 21’ 38.449’’ N, 82° 16’ 40.803’’ W), where the association between *H. unicolor* hamlets and the butterflyfish *C. capistratus* had been previously observed by Puebla et al. (2007). Fish were collected while SCUBA diving by hook-and-line or with hand nets at depths of between 3 −8m and then kept briefly in 80cm x 80cm x 50cm outdoor aquaria with running seawater, before data collection.

We considered a non-mimic hamlet, the barred *H. puella*, and two model-mimic putative pairs: i. the butter hamlet *(H. unicolor)* and its model, the foureye butterflyfish *(Chaetodon capistratus);* ii. the black hamlet *(H. nigricans)* and its model the dusky damselfish *(Stegastes adustus)*. In addition, we collected the two most common hamlet prey encountered in hamlet stomach contents, in the Bocas del Toro populations (Puebla et al. 2007), the masked goby *(Coryphopterus personatus)* and a mysid shrimp *(Mysidium columbiae). C. personatus* is an epibenthic small goby occurring in large schools hovering above coral heads and feeding on plankton (Bohlke and Robins 1962). *Mysidium columbiae* are among the most abundant swarming planktonic crustaceans on shallow coral reefs in the Gulf of Mexico and the Caribbean, generally found in patchy aggregations over corals or among mangrove roots (Wittmann and Wirtz 2019).

### Spectral measurements

#### i. Underwater spectral irradiances

We characterized the underwater photic environment on the Punta Caracol shallow coral reefs (Bocas del Toro, Panama) where the hamlet morphs, their models and their prey were collected. We used an Ocean Optics USB2000 spectrometer connected to an Ocean Optics ZPK600 UV/VIS optical fiber and fitted at one end with a CC-3-UV cosine corrector. The probe was secured at the tip of the longer arm of a white 1m long L-shaped pole and directed by a scuba diver vertically upwards and downwards, and horizontally perpendicularly to the shoreline, towards and away from shore, and the two directions parallel to the shoreline. This was repeated at different depths (just below surface, 2.5 m, 5.0 m, 7.5 m) just above hamlet territories. All measurements were taken between 11.00 and 13.00 on a sunny clear day.

The spectral distribution of irradiance was characterized by calculating λP50, the wavelength that halves the total number of photons (Munz and McFarland 1973), and the breadth of the light spectrum by calculating the difference Δλ between λP_25_ and λP_75_ cumulative photon frequency, in the range 350-650nm, most relevant for visual functions. The depth and wavelength dependence of the downwelling irradiance can be expressed in terms of the diffuse attenuation coefficient, K_d_(λ) (Mobley, 1994):

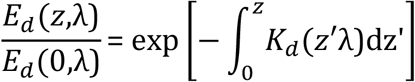

where *E_d_*(*z*,λ) is the downwelling irradiance at depth *z* and *E_d_*(0,λ) is the downwelling irradiance just below the water surface. If we consider the average 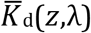 over the depth interval from below the surface to the maximum depth recorded, this is

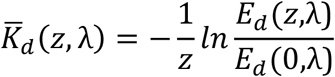

We calculated the average diffuse attenuation coefficient using the below-surface and 7.5 m depth downwelling spectral irradiance measurements.

#### ii. Fish body reflectance

A number of equivalent areas on the fish body showing clear differences in coloration between two or more species were identified and used to build a set of 7 ‘homologous’ landmarks, shared by all hamlets and their putative models. Reflectance measurements were taken from these landmarks across species, with a 200 μm UV/vis bifurcated fiber optic cable connected to an Ocean Optics USB2000 spectrometer and an Ocean Optics PX-2 pulsed xenon light source. Individuals were placed over a damp cloth and their skin maintained wet, following the guidelines of Marshall (2000). Reflectance records, obtained keeping both ends of the bifurcated fiber at approximately 45 degrees above the color patches, were calibrated against a Spectralon (Labsphere, North Sutton, NH) white standard.

### Receivers visual system

#### i. Masked goby visual sensitivity

We characterized the spectral sensitivity of the masked goby visual pigments by microspectrophotometry (MSP). Fish were maintained under dark conditions for a minimum of four hours prior to MSP and then euthanized with an overdose of MS-222 followed by cervical dislocation. The eyes were rapidly enucleated under dim red light, and the retinas removed and maintained in PBS (pH 7.2) with 6% sucrose. Small pieces of the retina were placed on a cover slide, fragmented to isolate individual photoreceptors, and sealed with a second cover slide and Corning High Vacuum grease. We used a single-beam, computer-controlled MSP, with a 100 W quartz iodine lamp that allowed for accurate absorption measurements down to 340 nm (Loew 1982; Losey et al. 2003). The peak of maximum absorption (λ_*max*_) of photoreceptors was obtained by fitting A1 or A2 templates to the smoothed, normalized absorbance spectra (Lipetz and Cronin 1988; Govardovskii et al. 2000). We used the criteria for data inclusion into the analysis of λ_*max*_ described in Loew (1994) and Losey et al. (2003).

#### ii. Masked goby lens transmittance

Lens transmission was measured directing light from the pulsed xenon light source through the lens mounted on a pinhole into a UV/vis fiber optic cable connected to the Ocean Optics USB2000 spectrometer. Lens transmission was expressed in terms of the 50% cutoff wavelength (*T_50_*) calculated from transmission spectra normalized to their maximum transmission between 300-700nm.

#### iii. Mysid shrimp visual sensitivity

Opossum shrimp (Crustacea: Mysida) have superposition eyes (Hallberg 1977). Visual sensitivity has been studied in depth in the genus *Mysis*, with evidence for a single visual pigment, with peak sensitivity positioned in the waveband 520-525nm in marine populations (Jokela-Määttä et al. 2005; Audzijonyte et al. 2012). In the absence of available data on visual sensitivities in *Mysidium*, we chose a value of 520nm, consistent with values in the related *Mysis*.

### Color and luminance discrimination

Vorobyev and Osorio’s (1998) color discrimination model was used to derive chromatic distances between corresponding color patches in hamlets, their models and natural backgrounds as perceived by a masked goby and by a mysid shrimp. Photoreceptor quantum catch Qi for a receptor of class i was calculated as:

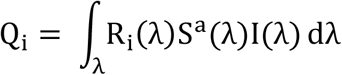

where R_i_(λ) is the absolute spectral sensitivity for receptor class i, S^a^(λ) is the reflectance spectrum of a color patch a, I(λ) is the irradiance spectrum, and integration is over the range λ = 350÷650 nm. Given the small distances (d=25-50cm) over which models and mimics interact with the receiver (i.e. the prey: masked goby or mysid shrimp) in clear coral reef waters, we considered light attenuation effects negligible.

If we take the signal fi of each receptor of class i as proportional to the natural logarithm of the receptor quantum catch qi, according to Weber-Fechner’s law, the contrast between two homologous patches, e.g. on the model vs on its mimic, will be:

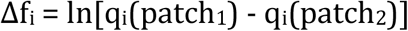

The Vorobyev-Osorio model assumes that discrimination thresholds are limited by photoreceptor noise. Color contrasts ΔS between corresponding patches on model and mimic viewed by the goby were calculated as:

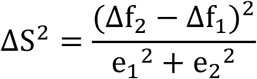

for a dichromatic species with short-wavelength (1) and long-wavelength (2) sensitive photoreceptors, with e_1_, e_2_, representing the photoreceptor noise associated with them. For a trichromatic species with short- (1), medium- (2) and long-wavelength (3) sensitive photoreceptors, the expression becomes:

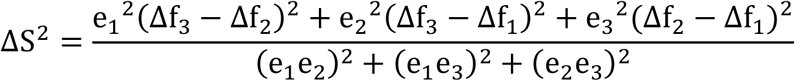

In relatively bright light conditions, as in the case of shallow waters in well-illuminated coral reefs, the photon shot component of noise is negligible and neural noise will be largely accounting for the photoreceptor noise ei. Neural noise is inversely proportional to the relative frequency of the receptor types as given by the following equation:

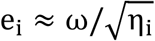

where ω is the Weber fraction and η_i_ the relative density of photoreceptors of type i. For the masked goby, we set a Weber fraction ω = 0.05 for the long wavelength sensitive cone, a conservative estimate for fishes (Cheney and Marshall 2009; Champ et al. 2016). During preliminary MSP work, we did not find any evidence of short-wavelength sensitive cones in this species. Taking a conservative approach, we decided to consider both a dichromat scenario with the MSP values obtained in this study as well as a hypothetical trichromat condition, adding a blue-sensitive cone located in a region of spectrum typical of other gobies (Table S1, Suppl. Mat.). For the dichromat, we set the relative proportions of the different cone types as 1:1 and at 1:4:4 for the trichromat scenario, to account for the apparent rarity (if present) of the putative short-wavelength sensitive cone class in this species. The Weber fraction for the mysid was set at 0.05.

Masked gobies and mysids might be able to discriminate mimic hamlets from models based on differences in luminance. Based on studies of other fish and terrestrial animals (Neumeyer et al. 1991; Kelber et al. 2003), we assumed that the longwave sensitive photoreceptor in the masked goby and mysid shrimp is responsible for the achromatic perception of luminance. Thus, we computed the luminance contrast ΔL between patches on model and mimic as:

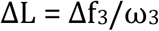

where ω = 0.05, a value close to that observed in other teleosts (Olsson et al. 2017). Color and luminance distances were calculated separately at three different depths, using horizontal irradiance (averaged over the 4 cardinal directions), measured at 2.5 m, 5.0 m and 7.5 m of depth, spanning the range of hamlet territories on the studied reef.

We tested whether perceptual distances in color (dS) and luminance (dL) of corresponding patches on hamlets and their putative models were statistically different with a PERMANOVA approach, using color and luminance distances in JNDs, with the *adonis* function in the R package *vegan* (Oksanen et al. 2007). Prior to this, the assumption of homogeneity of variances was tested for each patch across species with an analysis of variance followed by Tukey’s multiple comparisons tests. The distance-based PERMANOVA analysis was used to generate a pseudo-F statistics from the ratio of among/within distances between groups, and to obtain a null distribution by randomizing distances between observations (Anderson, 2005). We used 999 permutations to test for significant deviation from the null distribution and used the R^2^ as an estimate of effect size. Post-hoc tests were performed with the *pairwiseAdonis* function (Martinez Arbizu 2019), returning adjusted p-values on the pairwise comparisons.

We then assessed whether the effect size of the above differences between model and mimic was sufficiently large for the visual system of the receiver (i.e. the prey: masked goby, mysid shrimp) to perceive them. For each patch, we calculated distances in color space between the geometric means of each species and derived confidence intervals by applying a bootstrap procedure, as proposed by Maia and White (2018). All analyses were run separately using the mysid shrimp and the masked goby visual systems, the latter under both a dichromatic and a trichromatic scenario, and repeated at each of three depths.

### A prey’s view of natural scenes

We wished to gain insight into the spatial information available to masked gobies and mysid shrimp, when attempting the discrimination of a hamlet from its model in a natural scene. We applied the method of Caves and Johnsen (2018), implemented in the R package *AcuityView* (R development Core Team, 2016) which uses a Fourier transform approach to convert an image from spatial into frequency domain, then multiply it pixel by pixel by a modulation transfer function (MTF). This method uses Snyder’s (1977) MTF, which is a function of the minimum resolvable angle of the viewer. The result is an image that is devoid of all the frequencies above a threshold corresponding to the viewer’s acuity. After inverse Fourier transform to spatial domain, the image retains the level of detail that lies above the contrast threshold dictated by the viewer’s acuity and therefore provides us with insight into the amount of spatial resolution available to the prey viewing a scene including a predator, the hamlet, or a harmless ‘passer-by’, the model species.

Optical properties of the eye and spacing of photoreceptors and retinal ganglion cells are considered the main anatomical factors limiting acuity, with the former generally playing a minor role in affecting visual resolution in fishes. While photoreceptor densities are intuitively expected to influence the minimum resolvable angle (Northmore and Dvorak, 1979), visual processing by neural cells in the retina and, in particular, visual summation by ganglion cells, can significantly reduce resolution (in favor of increased sensitivity). Higher-level processing might, in certain cases, lead to further loss of spatial information (Warrant 1999). This is consistent with the frequent observation of higher anatomical than behavioral acuity values in fishes.

#### i. Masked goby acuity

We assessed the acuity of the masked goby *C. personatus* in terms of its optical anatomy, based on ganglion cell densities. Therefore, we consider these estimates as representing an upper limit to the masked goby acuity. To do this, we analyzed the population of ganglion cell layer (GCL) cells and estimated the upper limits of spatial resolving power from two retinas of *C. personatus*. The eyes were enucleated and fixed in paraformaldehyde (PFA) 4% diluted in phosphate buffer saline (PBS) for 3 hours, and then transferred to PBS and stored at 4º C. The cornea and lens were removed and the retinas were dissected from the sclera. The free-floating retinas were immersed in 10% hydrogen peroxide solution diluted in PBS, for 72 hours, at room temperature, for retinal epithelium bleaching. The retinas were washed in PBS and flattened onto a gelatinized glass slides with the ganglion cell layer facing up. The slides were exposed to 4% PFA vapors overnight, at room temperature, in order to increase the adhesion of the retina to the slide and for differentiation of the stained neurons (Stone 1981; Coimbra et al. 2006). For Nissl staining, the tissues were rehydrated by passing through ethanol series in decreasing concentrations (95%, 70% and 50%) and distilled water acidified with glacial acetic acid. The retinas were stained in an aqueous solution of 0.1% cresyl violet at room temperature for approximately 3 minutes and dehydrated by passing through a series of ethanol and xylene. The slides were coverslipped using DPX (Sigma-Aldrich, St. Louis, MO, USA). To evaluate shrinkage during the dehydration process, the areas of the retinas were measured before and after staining, using the software ImageJ (NIH, Bethesda, USA). Photographs of the retinas were taken with a digital camera (Axio CamMR, Carl ZeissVision GmbH, Germany), coupled to a stereomicroscope (SMZ775-T, NIKON, Japan), and software (Axio Vision 4.1, Carl Zeiss, Germany).

To estimate the total population of GCL cells, we applied the optical fractionator method (West et al. 1991) with modifications for retinal wholemounts (Coimbra et al. 2009, 2012). The retina was considered a single section, and so the section sampling fraction was equal to one. The total population of cells N_tot_ was estimated based on the total number of counted cells ΣQ and the area of the sampling fraction asf, which corresponds to the ratio between the counting frame and the sampling grid (Coimbra et al. 2009):

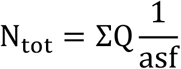

The cells were counted using a Leica DM5500B trinocular microscope with a motorized stage, connected to a computer running the Stereo Investigator software (MicroBrightField, Colchester, VT). The edges of the retina and the optic nerve were delineated using a 5x/NA 0.15 objective. Counts were made using x100/NA 1.4-0.7 oil immersion objective, at regular intervals defined by a sampling grid at 150×150 μm placed in a random, uniform and systematic fashion, covering the entire retinal area. An unbiased counting frame at 50×50 μm was imposed at each sampling frame and cells were counted if they lay entirely within the counting frame or if they touched the acceptance lines without touching the rejection lines (Gundersen 1977). The coefficient of error was calculated using the method of Scheaffer et al. (1996). All cellular elements located within the GCL were counted, independent of size (Collin and Pettigrew 1988a, 1988b; Collin 1989; Collin and Partridge 1996).

The theoretical spatial resolving power of each eye was estimated based on the maximum density of presumed retinal ganglion cells D and the focal length of the eye f, obtained from a Matthiessen’s ratio of 2.34, appropriate for gobies (Matthiessen 1880; Hansen 1988; Wanzenbock et al. 1996; Ota et al. 1999). Considering a square array of retinal ganglion cells, the mean cell-to-cell spacing S is related to the maximum density D of ganglion cells per mm^2^ through the expression S^2^ = 1/D. The maximum spatial (Nyquist) frequency (ν) of a sinusoidal grating resolvable by a square arrangement is ν = 1/(2ΔΦ), where ΔΦ is the inter-receptor angle ΔΦ = S/f (Snyder and Miller 1977). Substituting, we obtain:

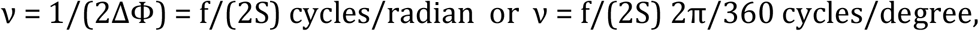

or its inverse, the smallest resolvable angle α, in degrees.

#### ii. Mysidium shrimp acuity

Behavioral estimates of visual acuity are available for the mysid shrimp *Mysidium columbiae*, based on optomotor response experiments (Buskey 2000). Under conditions of optimal illumination, individuals placed in an optokinetic drum, followed vertical black and white stripes of varying width consistently resolving differences down to 6mm from a minimum distance of 15mm. Given the known relationship between subtended angle α, reactive distance d and stripe width w: α = 2arctg(0.5w/d), this corresponds to a subtending arc of 7.66 degrees, the value of behavioral acuity we used in the spatial patterns analyses.

#### iii. image collection

Underwater scenes were captured with either a Canon 5D Mark III camera with a 24-70mm f2.8 lens set on 50mm or with a Canon G7X. Two sets of high-resolution photos were taken. First, hamlets and their models were photographed underwater at a depth of about 5m, from an approximately horizontal line of view, against different natural backgrounds; ii. In addition, during preliminary observations, we noted that the ventral silhouette of the butter hamlet *H. unicolor* closely resembles in shape and colors that of its putative model, the butterflyfish *C. capistratus*, despite having very different lateral profiles. In order to compare the appearance of the butter hamlet and its model (and, for comparison, the black hamlet *H. nigricans* and the barred hamlet *H. puella*), as seen from below at a 45° angle, as a masked goby hovering above a coral is likely to do (see also next paragraph), we collected one individual for each hamlet morph and a four-eye butterflyfish. Fish were euthanized with an overdose of MS-222 in an aerated aquarium then moved for few minutes at −20°C. A hole was then drilled on the dorsal area of each fish allowing a thin metal bar to be passed through the body so that the butter, black, and barred hamlets and the four-eye butterflyfish could be placed side by side on a common base and brought underwater. A series of photos were taken at a depth of 5m in proximity to hamlets territories, at about 45° and 40cm from the metal rod placed against different natural backgrounds. We did not note appreciable differences in color patterns between live non-stressed individuals as observed in the wild and the sacrificed individuals as treated in our protocol, but their eyes did take a cloudy appearance and the tips of their fins showed some minor damage. However, it is significant that once we completed photographing and the bar was left for few minutes above a dead coral head, two hamlets, a blue and a butter, came immediately to inspect our setup and vigorously displayed against the “intruding” hamlets on the bar without interrupting their territorial displays even when we approached to try and recover the bar.

#### iv. image processing and analysis

Following Caves and Johnsen (2018)’s guidelines, high resolution photos were cropped to 1024×1024 pixels and saved separately for each RGB channel. The angular width of each scene was obtained by scaling with an object of known actual size in the photo and by selecting a biologically significant viewing distance. For the photos of live models and mimics in their natural habitat (*i*. above), we measured the sizes of corals or rocks appearing in the images, while for the bottom-up images of the mounted individuals (ii., above) we measured the hamlets’ inter-orbital distances, to derive image width.

Underwater observations of feeding hamlets in Bocas del Toro populations (MERP, *pers. obs*., 2015-2017) showed that they can rush from a distance of more than 4ft towards a dense cloud of mysids or masked gobies, stopping for about 1-2 seconds at a distance of less than 50cm (i.e. about five body lengths) away from the prey, for precision aiming at a single individual. They then strike horizontally when preying on mysids and from above at about 45° when striking at masked gobies. Given this pattern of predatory behavior, we considered that a mysid or a masked goby would have a reasonable chance of evading a predatory strike if it recognized an approaching predator and initiated escape at a distance over three hamlet body lengths away, i.e. about 25cm distance. We used these two values, 50cm and 25 cm as the relevant viewing distances d, in the image analysis. The angle α subtending the scene is then α = 2arctg(actual width/2d).

## RESULTS

### Spectral measurements

#### i. Underwater spectral irradiances

Downwelling spectral irradiance over our study site had typical characteristics of tropical reef waters (Figure 1) with similar λP_50_ across depths (under-the-surface: 516nm; 2.5 m: 516nm; 5.0 m: 512nm; 7.5 m: 520nm) and values very close to those measured by McFarland and Munz (1975) on a Pacific atoll (λP_50_ = 518nm at 5m depth). In contrast to that study, however, we observed a more pronounced effect of attenuation on the irradiance spectrum, particularly at longer wavelengths. The average diffuse attenuation coefficient of the downwelling irradiance was highest at 680nm (K_d_ = 0.143 m^-1^) with a second maximum in the near-UV (K_d_ = 0.106 m^-1^). The minimum was attained at 350nm, the shortest wavelength recorded (K_d_ = m^-1^) and a second minimum at 530nm (K_d_ = 0.044 m^-1^) (Figure S1, Suppl. Mat.). This pattern of higher attenuation with depth at short and long wavelengths is mirrored by the narrowing of spectral bandwidth (Δλ) with depth (Figure 1, main text; Figure S1, Suppl. Mat.). In clear atoll waters, at 5m depth, McFarland and Munz (1975) reported a Δλ = 105nm, while at Punta Caracol, in Bocas del Toro, at a similar depth (5.0 m) we measured a substantially narrower bandwidth (Δλ = 75nm), a value observed only at 20 m in their study. Upwelling irradiance at maximum depth, closer to the bottom of this shallow reef system, was shifted to longer wavelength (Figure 1) as a result of the mixed composition of the bottom, consisting of coral outcrops interspersed with yellow sand patches. As the distance from the bottom increased, the up-welling spectrum shifts to shorter wavelengths and to a spectral profile similar to the down-welling and side-welling spectra.

**Figure 1.**
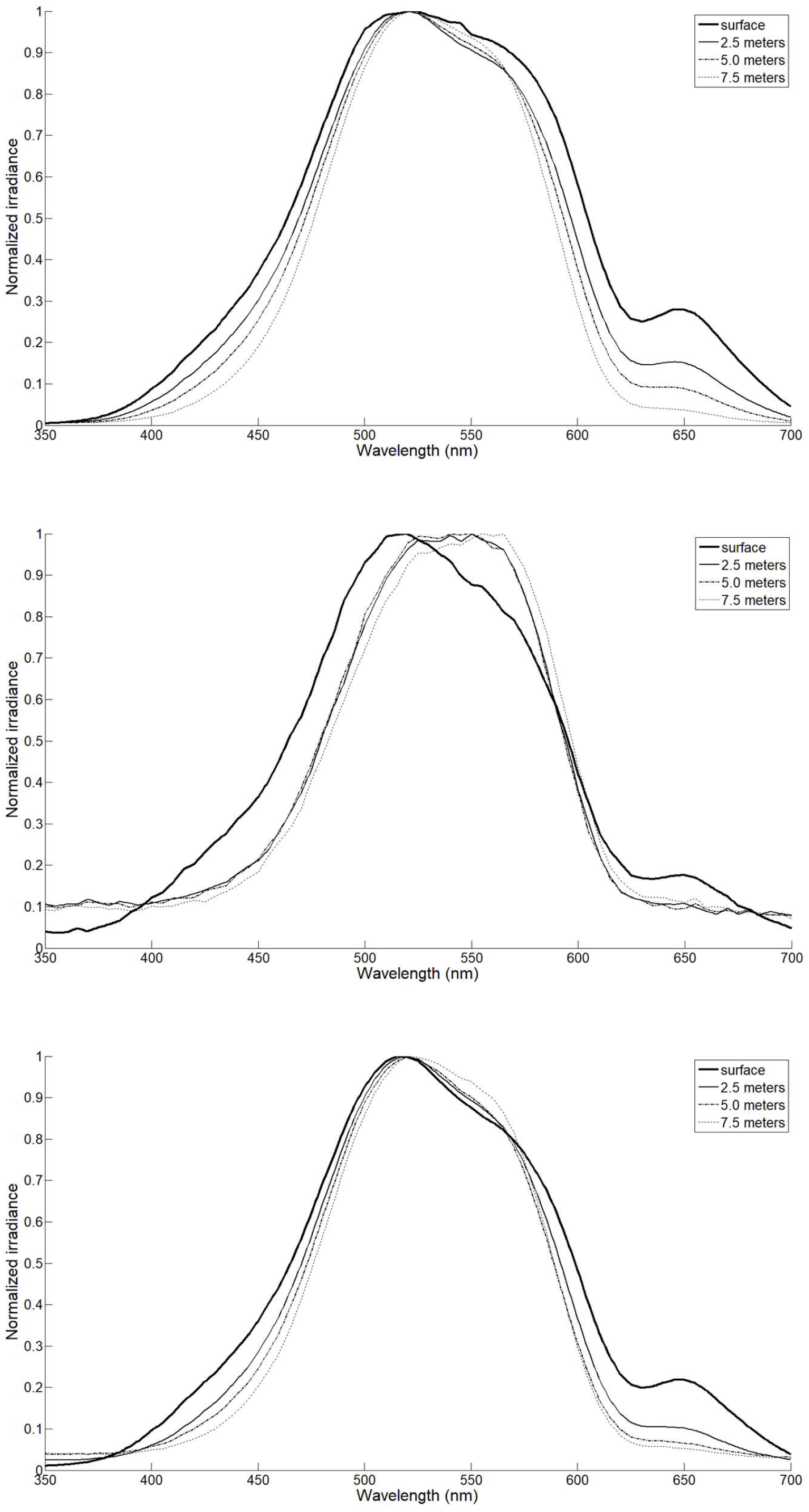
Down-welling (*top*), up-welling (*center*), and average side-welling (*bottom*) irradiances, measured at Punta Caracol, Bocas Del Toro, Panama, on a vertical depth profile, just under the surface (—), 2.5m (—), 5.0m (---) and 7.5m (…).

#### ii. Fish body reflectance

Spectral reflectance measurements were collected from 7 homologous landmarks on the body of 42 individuals across 5 species. Mean reflectance spectrum and confidence intervals for a representative spot (pelvic fin) are shown in figure 2.

**Figure 2.**
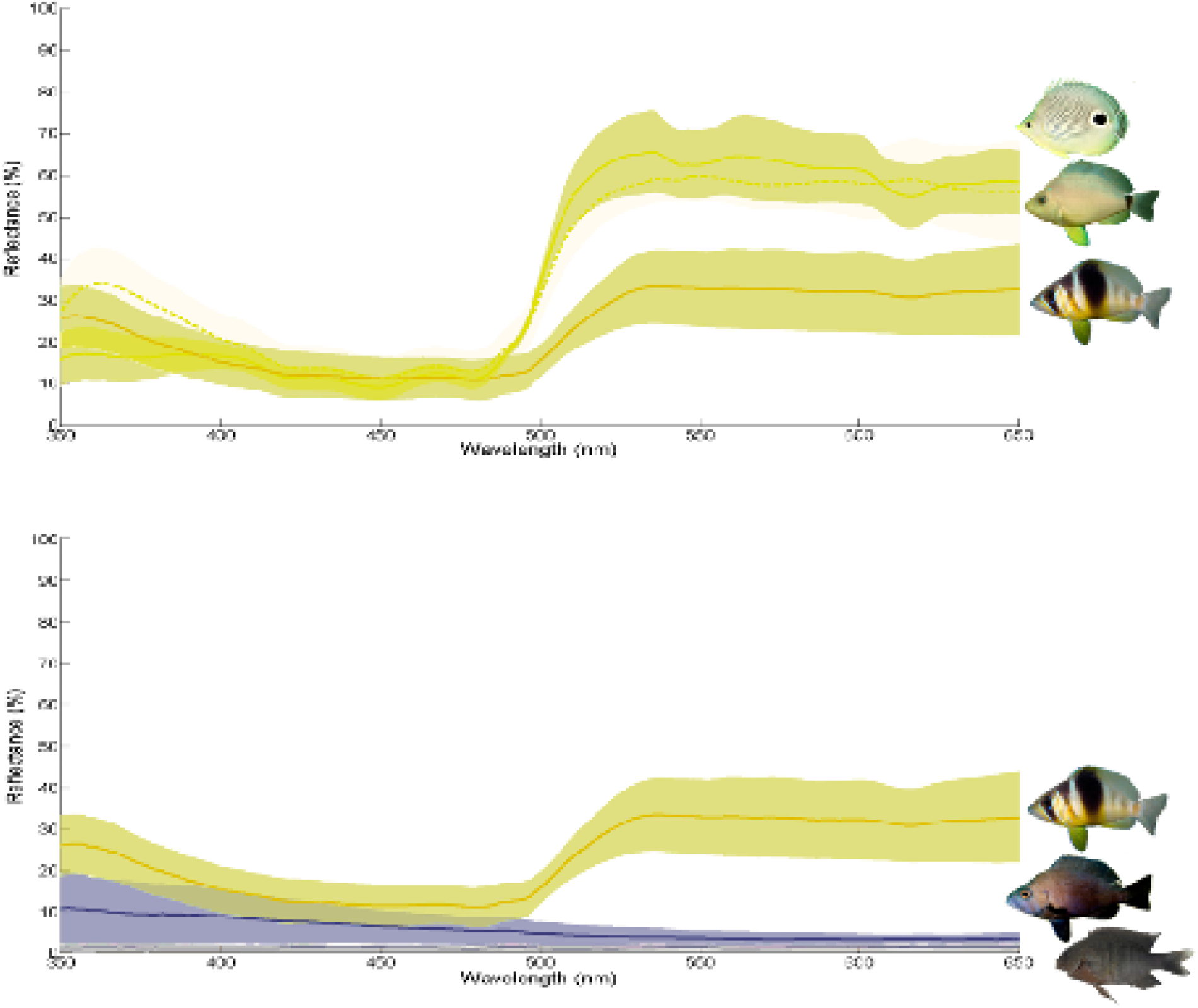
Mean spectral reflectance and confidence regions of body patch #5 (center of left pelvic fin) across species. *Left:* from top, four-eye butterflyfish *C. capistratus* (putative model), butter hamlet *H. unicolor* (putative mimic) and the non-mimic barred hamlet *H. puella. Right:* from top, the non-mimic barred hamlet *H. puella)* and the black hamlet *H. nigricans* (putative mimic), and bottom, the dusky damselfish *S. adustus* (putative model). In the inset, the location of the final seven patches used in the analyses.

### Receivers visual system

#### i. Masked goby visual sensitivity

We collected spectral sensitivity data from the retinas of n = 4 *C. personatus* individuals. The retina of masked gobies appeared rod-dominated (rhodopsin λ_max_ = 500nm). Double cones showed either the same visual pigment with λ_max_ = 539nm or one member with λ_max_ = 531nm and the other with λ_max_ = 539nm. We did not find any evidence for short wavelength cones. The very small distance between the two green-sensitive cone λ_max_ (531nm, 539nm) is unlikely to provide the masked goby with true color vision but might confer the fish broader sensitivity in that particular region of the light spectrum. Overall, these results are in line with previously reported spectral sensitivities from other tropical gobies (Table S1). In particular, the absence of a short wavelength cone in *C. personatus* is a condition shared with the only other coral reef goby for which MSP data are available. However, we cannot exclude that small blue or violet cones might be present in very low frequencies in the masked goby retina, given the sampling design typical of micro-spectrophotometry. For this reason (see below), we considered both a visual model that includes the cone repertoire found by our MSP study and an additional one that incorporates a third short wavelength cone, positioned in the region of sensitivity characteristic of other gobies (λ_max_ ≈ 455nm), that would potentially confer trichromatic color vision to the goby (Fig. 3 *left*, main text; Table S1, Suppl. Mat.).

#### ii. Masked goby lens transmittance

We examined the eye lenses of n = 3 individuals. The wavelength at which 50% of the maximal transmittance was reached (T_50_ = 410÷411nm) and the shape of the transmittance curve suggest that there is little scope for UV signals reception in this goby species (Figure 3, *right*).

**Figure 3.**
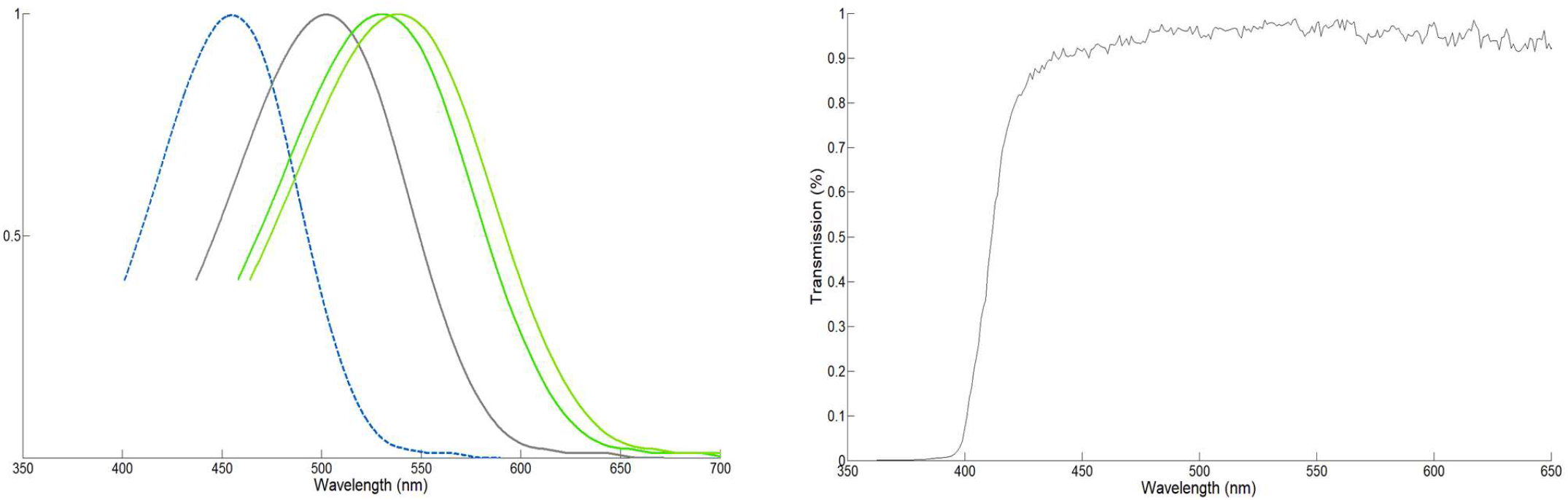
*Left:* Absorbance templates (from Lipetz and Cronin 1988; Govardovskji et al. 2000), representing the rod (501nm; in grey) and cone visual pigments (521nm, 539nm; in green) found in the retina of the goby *Coryphopterus personatus;* an additional blue cone pigment (455nm; in blue) was added, in an alternative modelling scenario (see main text). *Right:* Lens transmission spectrum of the masked goby *C. personatus*. The T50 is located at 410÷411 nm.

### Color and luminance discrimination

#### i. Masked goby

The PERMANOVAs of differences in dS and dL between corresponding patches across species, calculated at different depths, and with either two (531nm, 539nm) or three (455nm, 531nm, 539nm) cone classes, were all significant (p < 0.001; Table S2, Suppl. Mat.). Post-hoc tests on chromatic distances dS showed that for almost all species pair contrasts, at least three or more patches on the body were significantly different between species. The exceptions were the two putative model-mimic pairs: the butter hamlet *H. unicolor* and its model, the butterflyfish *C. capistratus*, were not significantly distinguishable in color at any of the measured patches, when the viewer had the 531/539nm cone set and distinguishable by one patch (#1) only, when the viewer was provided with a 455/531/539nm cone set, and this irrespective of depth. In the black hamlet *H. nigricans* and its putative model, the damselfish *S. adustus*, two patches (#3, #7) were significantly different in color between species when seen by a 531/539nm viewer and three patches (#1, #3, #7) when seen by a 455/531/539nm viewer, although not at all three depths (Table S2, Suppl. Mat.). Post-hoc tests on achromatic distances dL reveal that at least four and up to all seven patches were significantly different (p < 0.05) across species at any depth, with the only exception of the model-mimic pair butter hamlet *H. unicolor* and butterflyfish *C. capistratus* for which all seven patches were indistinguishable (p > 0.21) between species, irrespective of depth and cone set of the goby.

Whether the significant differences in color and luminance found between species are of a magnitude detectable by the viewer was tested by calculating distances between the geometric means of each species and generating confidence intervals by bootstrapping (Maia and White 2018). We found that the only patch that was significantly different in the PERMANOVA analysis between the *H. unicolor – C. capistratus* mimic-model pair, when the goby was provided with three visual pigments 455/531/539, is effectively indistinguishable by the goby (dS < 0.31) at all depths, as are all other patches (Figure 4), and this holds true irrespective of depth and cone pigment repertoire of the goby (Figure 5 and Figure S2, Table S3, Suppl. Mat.). This result suggests that the *H. unicolor – C. capistratus* pair fulfills the requirement of a model-mimic relationship, in terms of color differences since color distances between corresponding patches on model and mimic are well below the discrimination threshold of the signal receiver, the goby. In the other putative mimic-model pair, *H. nigricans* – *S. adustus*, the various patches that we found significantly different in the PERMANOVA analysis, had color distances of magnitude below or at the discrimination threshold (Figure 4), and that holds true at all depths and cone set conditions (Table S3, Suppl. Mat.), in line with a model-mimic hypothesis. All other species pair contrasts were above threshold when modelled with a 455/531/539 cone set, while chromatic contrasts between the pair *S. adustus – C. capistratus* were below threshold when modelled with a 531/539 goby visual system (Table S3, Suppl. Mat.).

**Figure 4.**
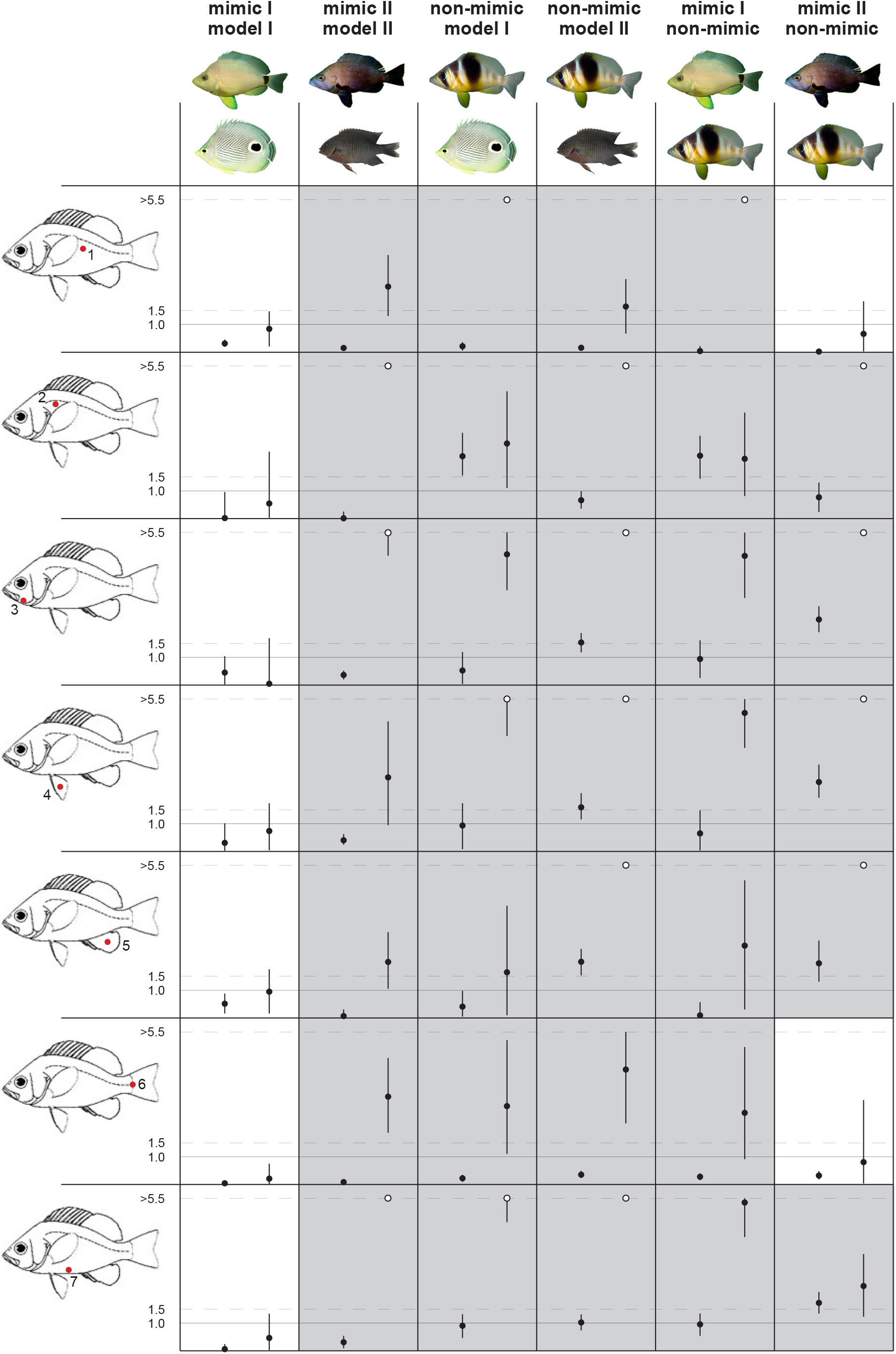
Color distances dS (on the left, in each box) and achromatic distances dL (on the right, in each box) and their bootstrap confidence limits, for each species pair viewed by the prey, a masked goby visual system with 455nm, 531nm, 539nm cone pigments, at a depth of 5m. A continuous line, corresponding to dS (or dL) = 1, marks the perceptual threshold, below which a particular patch (#1-7,indicated in the sketch on the left) is likely indistinguishable by the viewer. Distances dS, dL >5.5 are presented with an open dot at the top end of their respective box. Discriminable patches (either because dS or dL >1, or both) are represented by grey-shaded boxes.

The bootstrap analysis of achromatic contrasts revealed that only one species pair is indistinguishable by the goby and that is the putative mimic-model pair *H. unicolor – C. capistratus* (mean dL < 1 for all patches: Figures 4, 5), consistent with a model -mimic scenario. All other species contrasts contain at least four patches that can be discriminated between species by the goby based on achromatic distances, including the second putative mimic – model pair, *H. nigricans – S. adustus*, for which all seven patches can be discriminated by the goby (Figure 4 and Figure S2, Table S3, Suppl. Mat.). This suggests that *H. nigricans* and *S. adustus* do not represent a model – mimic pair, at least for the goby *C. personatus*.

**Figure 5.**
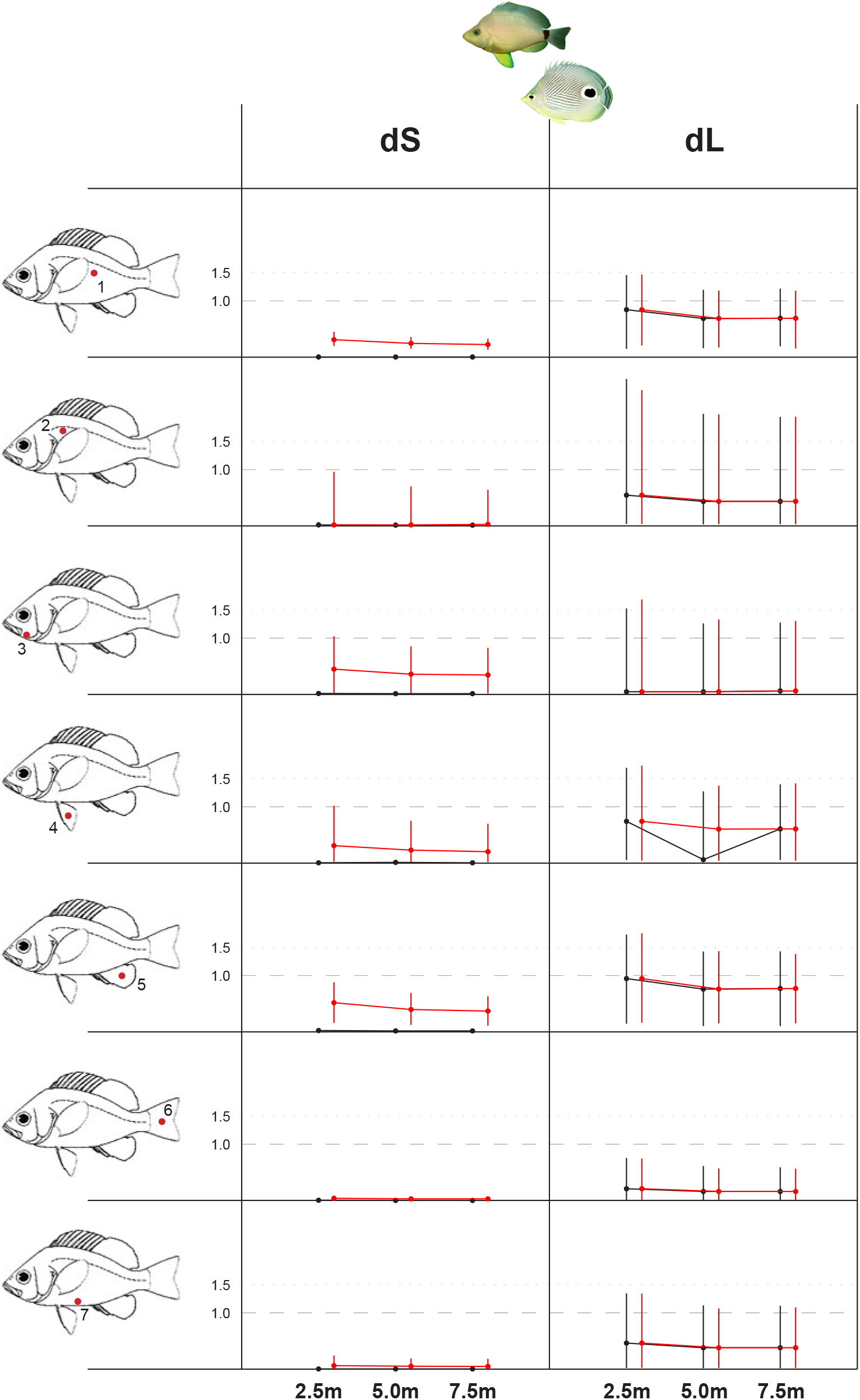
Chromatic (dS) and achromatic distances (dL) and their bootstrap confidence limits, for the mimic-model pair *H. unicolor* and *C. capistratus*, at different depths (2.5m, 5.0m, 7.5m), viewed by the prey, a masked goby visual system with 455nm, 531nm, 539nm cone pigments.

#### ii. mysid shrimp

The PERMANOVAs of differences in dL between corresponding patches across species, modelled with a single visual pigment and calculated at different depths were all significant (p < 0.004). Post-hoc tests showed that at least four and up to all seven patches were significantly different (p < 0.05) across species at any depth, with the single exception of the model-mimic pair butter hamlet *H. unicolor* and butterflyfish *C. capistratus* for which all seven patches were indistinguishable (p > 0.23) between species, irrespective of depth (Table S4, Suppl. Mat.).

The bootstraps of species groups showed that at least five patches were distinguishable (i.e. above detection threshold dL > 1) in any two species contrast (Figure S3, Table S5, Suppl. Mat.). The notable exception is the pair *H. unicolor* - *C. capistratus*, for which all spots were below detection (dL < 1), suggesting that putative mimic *(H. unicolor)* and model (*C. capistratus*) are not discriminable by the mysid shrimp based on achromatic contrast between corresponding patches. The achromatic differences in the second putative pair, *H. nigricans* - *S. adustus*, are all above threshold.

### A prey’s view of natural scenes

Visual acuity in the masked goby was estimated from the total number of cells in the ganglion cell layer of two retinal wholemounts from different *C. personatus* individuals. An average cell population of 179,769 ± 12,914 was obtained (Table 1), with high densities observed in the retinal periphery and peak density located in the ventral region (89,400 cells/mm^2^) while lowest densities were observed in the central retina (1,200 cells/mm^2^). The upper limit of the spatial resolving power estimated from lens radius and the maximum density of GCL cells was 2.356 ± 0.14 corresponding to a minimum resolvable angle (MRA) α = 0.425 ± 0.02 (Table 1). The number of sites counted for each retina, Scheaffer’s coefficient of error (CE) and area of sampling fraction (asf) are described in Table 1. Shrinkage was below 5% and considered negligible (Coimbra et al. 2006). For the mysid shrimp, we used Buskey (2000)’s acuity estimate of α = 7.66 degrees (or 0.13 cycles/degree).

**Table 1.**
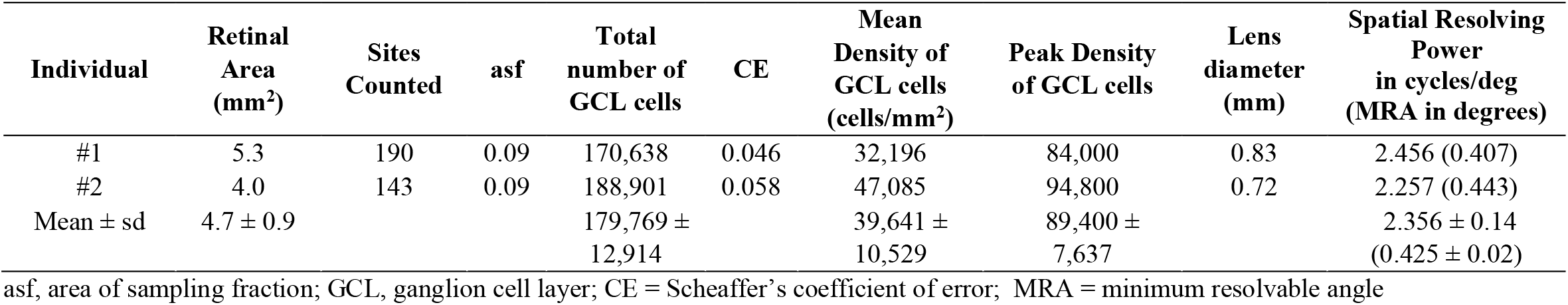
Stereological assessment of the population of cells in the retinal GCL of *Coryphopterus personatus* and the anatomical parameters used to estimate the upper limit of spatial resolution.

| Individual | Retinal Area (mm<sup>2</sup>) | Sites Counted | asf | Total number of GCL cells | CE | Mean Density of GCL cells (cells/mm<sup>2</sup>) | Peak Density of GCL cells | Lens diameter (mm) | Spatial Resolving Power in cycles/deg (MRA in degrees) |
| --- | --- | --- | --- | --- | --- | --- | --- | --- | --- |
| #1 | 5.3 | 190 | 0.09 | 170,638 | 0.046 | 32,196 | 84,000 | 0.83 | 2.456 (0.407) |
| #2 | 4.0 | 143 | 0.09 | 188,901 | 0.058 | 47,085 | 94,800 | 0.72 | 2.257 (0.443) |
| Mean ± sd | 4.7 ± 0.9 |  |  | 179,769 ± 12,914 |  | 39,641 ± 10,529 | 89,400 ± 7,637 |  | 2.356 ± 0.14 (0.425 ± 0.02) |

asf, area of sampling fraction; GCL, ganglion cell layer; CE = Scheaffer's coefficient of error; MRA = minimum resolvable angle

The underwater images taken at Punta Caracol and Punta Juan, in Bocas del Toro, Panama, were modified to account for the spatial resolution of *C. personatus* masked gobies and *Mysidium* shrimp. After processing, they provide a first approximation of scenes including models or putative aggressive mimics, as perceived by the prey, masked goby or mysid shrimp, given their visual acuity. The Fourier analysis of natural scenes suggests that *Mysidium columbiae* shrimp are only able to discern, even at relatively small distances (25cm) bright moving vs dark moving objects in their field of view. While temporal resolution was not considered in this study, overall the visual system of this mysid shrimp would be unable to distinguish, at the distances considered, a harmless moving target from a predator, based on visual cues, with the exception, possibly, of information regarding the target’s direction. The visual acuity of masked gobies *C. personatus*, in contrast, was sufficient to gather relevant information about other species’ general features at both distances, despite poor color vision.

The juxtaposition of model and mimic *H. unicolor-C. capistratus* in figure 6, as seen by a masked goby, reveals how the deceit might be obtained. The most relevant features shared by both model and mimic (figure 6, second row, left) are a uniform bright (yellow) body coloration with no vertical barring, a large black spot at the base of the tail highlighted by a white ring, the yellow tip of the snout, and the bright yellow pelvic fin. In addition, moving in a three-dimensional space, in the eyes of a receiver with little or no depth perception, will make the silhouette of the targets change continuously, making overall shape differences between model and mimic an unreliable cue for species discrimination.

**Figure 6.**
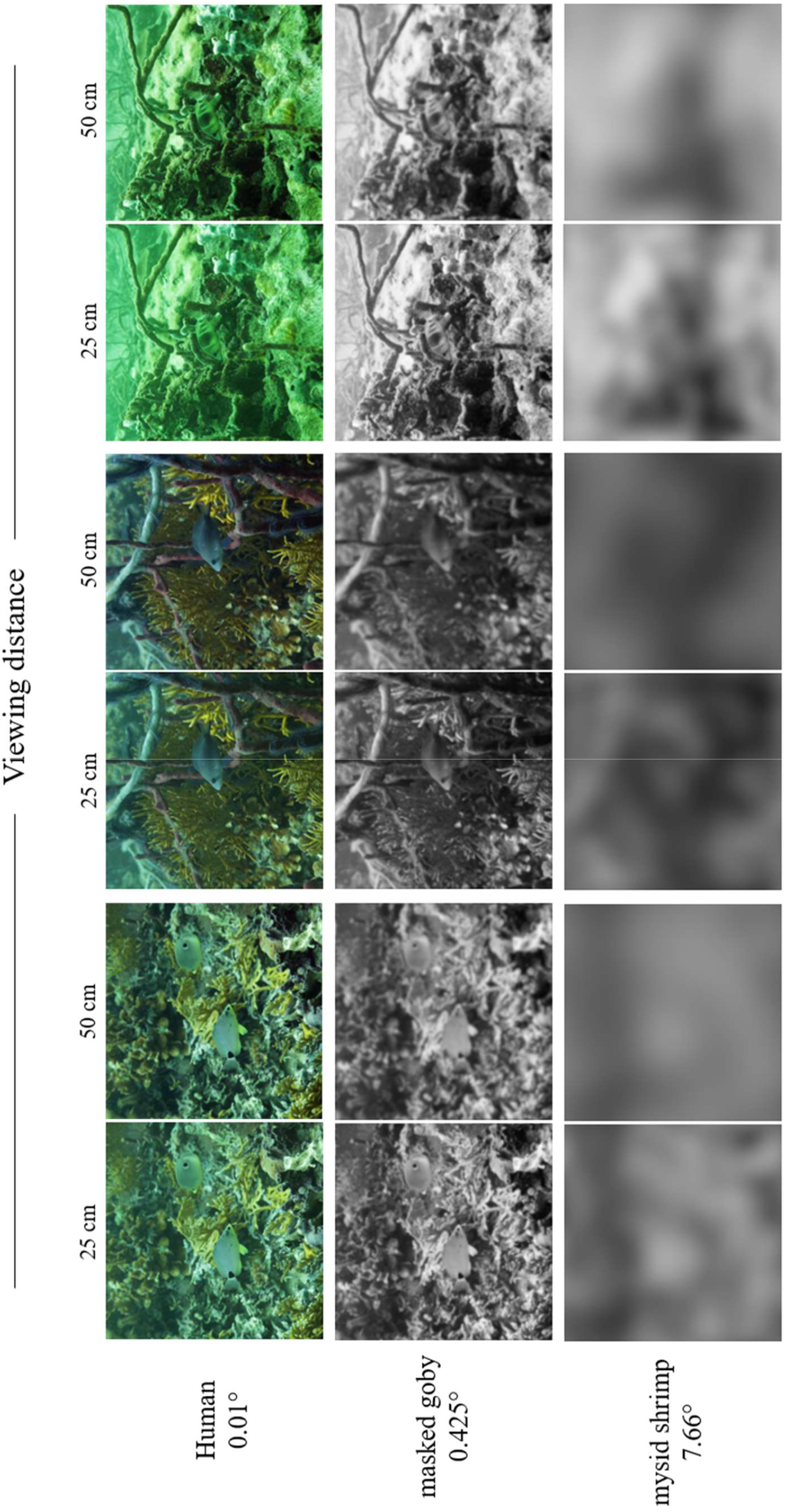
Images of *Hypoplectrus* hamlets in a natural scene, at a depth of 5 meters, on a coral reef in Bocas del Toro, Panama, as seen by humans and by the two main prey of hamlets, i.e. the masked goby *C. personatus* and a *M. columbiae* shrimp, at the distances in which interactions between these species are known to occur. First two columns, on the left, butter hamlet *H. unicolor* and its model, the foureye butterflyfish *C. capistratus*. Central columns, the black hamlet *H. nigricans*. Last two columns, on the right, barred hamlet *H. puella*. Acuity, expressed as minimum resolvable angle in degrees, is shown for each viewer.

The black hamlet *H. nigricans* is likely to gain an advantage in camouflage when moving between corals in sheltered areas where it will be best protected from predation and inconspicuous to its prey. However, its homogeneously dark coloration will stand out against open water and brightly lit corals and sponges and sand, making it a very conspicuous threat in the eyes of both masked goby and mysid shrimp. This suggests a defensive origin to this color morph. The barred hamlet *H. puella* (Figure 6, two columns on right-end side) sports dark vertical bars that are certainly a potential cue of a predator closing in, for a masked goby. A pattern of light background with vertical bars is shared with other potential predators of the masked goby so it would be adaptive to flee whenever an object with vertical barring is approaching. Indeed, we believe the concealment of vertical barring in the butter hamlet *H. unicolor* was the single most significant phenotypic change for the origin of the efficient deceit on masked gobies. The images also show that *H. puella’s* colors and vertical bars act as extremely efficient camouflage by disruptive patterning, for all three visual systems, including our own, another case of defensive origin for a hamlet color morph.

## DISCUSSION

The extraordinary variation in color patterns between *Hypoplectrus* hamlets has long been attributed to aggressive mimicry (Randall and Randall 1960; Thresher 1978; Fischer 1980; Domeier 1994; Puebla et al. 2007). However, apart from such comparisons frequently based on color flash-enriched photos, no attempt has been made to examine the resemblance of hamlets to the proposed models from the meaningful perspective of the intended signal receiver under natural light conditions. In a case of aggressive mimicry such receiver is a hamlet prey. Here we examined two putative mimicry pairs in the *Hypoplectrus* hamlet complex as well as a non-mimic hamlet, from the point of view of the visual system of two ecologically and taxonomically distinct prey species, the masked goby *C. personatus*, and the planktonic mysid shrimp, *M. columbiae*. We evaluated the prey’s discriminating ability in terms of differences in hue, luminance and color pattern of models and mimics and considered the effects of depth on the efficacy of mimicry. We found that one putative model-mimic pair, the butter hamlet and the foureye butterflyfish, are the only pair-wise species comparison that is indistinguishable in the eyes of the prey, while all others considered are above the threshold of discriminability. This, together with behavioral evidence for the association of this hamlet morph with its model butterflyfish (Puebla et al. 2007; Picq et al. 2019), constitutes strong support for an aggressive mimicry scenario in this pair. A second putative model-mimic relationship *(H. nigricans* black hamlet - S. *adustus* dusky damselfish), on the other hand, is not supported by our data, since discrimination between the two is well above perceptual threshold by the visual systems of both masked goby and mysid shrimp.

Different visual cues (color, luminance, dark barring) are relevant for the discrimination of different hamlet morphs from their harmless models by either prey. However, depth had very limited influence on visual thresholds, not surprisingly given the narrow depth range in which the majority of hamlet territories were located in our study population. In both chromatic models, the ability of the masked goby to discriminate the predator butter hamlet *(H. unicolor)* from its putative model, the four-eye butterflyfish (*C. capistratus)* was well below threshold both in terms of color and luminance, suggesting the prey would not be able to separate model from mimic based on differences in hue or brightness, regardless of depth.

Modeling a mysid shrimp’s visual system, which is devoid of color perception, revealed that only butter hamlets and their putative model butterflyfish could represent a valid model-mimic pair, the difference in achromatic contrast between the two being well below the threshold of a mysid discrimination ability. On the contrary, the shrimp’s visual system has the potential to discriminate efficiently all other putative pairs on the basis of achromatic contrast. In conclusion, at least for the masked goby *C. personatus* and the mysid shrimp *M. columbiae*, known to be the principal diet items in the Bocas del Toro hamlet populations, the butter hamlet *(H. unicolor)* represents the only case consistent with an aggressive mimicry scenario, if color and/or luminance are used by the prey to discriminate friend (the model) from foe (the mimic). In addition, the analysis of color patterns of model and putative mimic in a natural scene, at biologically realistic distances, based on the visual acuities of both preys, suggests that differences in the fine patterns over a relatively homogeneous yellow coloration of butter hamlet and butterflyfish are imperceptible.

An alternative strategy seems to have been taken by the black hamlet *(H. nigricans)*. Its uniformly blue-black coloration did not fulfil the criterion of resemblance to a dusky damselfish (*S. adustus*) model, as both masked goby and mysid shrimp are likely to discriminate with efficiency the two, on the basis of color and/or luminance. However, particularly in the relatively turbid underwater conditions in Bocas del Toro, black hamlets can be quite hard to spot when not swimming well above the reef, and this is even more the case for the generally limited spatial resolution of its prey. While it is unclear whether this dark cryptic coloration confers any advantage to the black hamlet in approaching its prey, it likely provides this morph some protection from its predators, as it does the dark brown cryptic coloration of its hypothetical model, the dusky damselfish *S. adustus*.

The non-mimic barred hamlet *(H. puella)* was clearly discriminated from other hamlets and putative models. Even when considering the most similar color patches between butter and barred hamlet, masked gobies and mysid shrimp appear to be always able to discriminate between the two hamlet morphs (while they cannot between butter hamlet and its model butterflyfish), suggesting that the butter hamlet divergence from the barred morph might indeed confer an advantage in approaching a prey over the barred form. In addition, a barred hamlet at close distance from its prey can be easily identified by the typical highly contrasting dark vertical bars over a comparatively brighter yellowish body, even by the very limited acuity of a mysid, as suggested by the Fourier transform analysis. Interestingly, the bright areas of the barred hamlet body are very close in color to a number of reef substrates we measured (Pierotti et al. in prep), suggesting that, in combination with the dark body areas, this might grant good camouflage by disruptive coloration, protecting the hamlet from its own predators. This in turn might lead to a trade-off between predator avoidance by the barred hamlet at the expense of its own predatory efficiency, by increased conspicuousness at short range when approaching a prey, particularly from above, against an open water background. It is interesting to note that barred and more complex disruptive patterns are also typical of other sympatric basses and all invariably attack their prey almost horizontally while close to the substrate. On the contrary, butter hamlets are often seen attacking from about 45 degrees above masked gobies (MERP, pers. obs.). From this line of view the hamlet silhouette and color is remarkably similar to that of a four-eye butterflyfish, in particular the very conspicuous structural-yellow pelvic fins (Figure S4, Suppl. Mat.). Notably, while the color of pelvic fins in hamlets is highly variable between individuals within morphs, from colorless to bright yellow to highly saturated indigo or blue, this does not seem to be the case for butter hamlets, invariably sporting bright structural yellow on their pelvics with very similar hue to that of four-eye butterflyfish pelvics, as evidenced by the reflectance measurements and below-threshold perceptual distances. Therefore, the observation that butter hamlets attack masked gobies from slightly above suggests they might be behaviorally optimizing the efficiency of their aggressive mimicry by presenting to the prey with best visual abilities, the masked goby, the closest model-resembling area of their body. We did not observe such top-down attack strategy by the butter hamlet when preying on mysids, possibly because of the mysid’s limited visual system.

In conclusion, this set of results, together with previous behavioral work by Puebla et al. (2007; 2018) and Picq et al. (2019), lends support to the aggressive mimicry scenario for the butter hamlet and its model, the four-eye butterflyfish, at least in the Bocas del Toro population, where the ‘model following’ behavior was observed and where this study was conducted.

Did the color patterns of the butter hamlet evolve for efficient aggressive mimicry by imitating in color and behavior a common and harmless (to their prey) butterflyfish or other selective forces brought about the similar appearance which was subsequently recruited for aggressive mimicry? Picq et al. (2019) found that the model-tracking aggressive mimicry behavior of butter hamlets represents in fact one of two alternative behavioral syndromes, associated with territoriality. Territory holders with defined permanent hide-outs only rarely engaged in aggressive mimicry, while roaming individuals, lacking a defined permanent hide-out and territory systematically took advantage of this behavioral strategy. The authors’ results suggest that the aggressive mimic strategy confers an advantage in terms of foraging opportunities, at the expense of higher exposure to predators and possibly fewer mating opportunities. While the factors affecting the quality of hamlets territories and/or hide-outs are unknown at present, it is reasonable to assume that typical hideouts with multiple entrances of limited size (MERP, pers. obs.) in the proximity of clusters of epibenthic mysid shrimp and/or *Coryphopterus* gobies would provide the benefits of foraging under low predation risk. The location of these prey clusters above particular reef spots is remarkably stable. A suitable safe hide-out near a foraging source is also likely to allow resource defense from competing conspecifics, especially roaming individuals. The alternative strategy of model-tracking aggressive mimics, while possibly providing an advantage in foraging efficiency inevitably comes at the cost of increased exposure to predators. We speculate that optimal hideouts and territories are likely to be in limited numbers over a reef leading some individuals to be out on the reef with no stable hideout and territory forced to move from a food patch to another, while exposed to conspecific aggression and to predation. In this context, before aggressive mimicry evolved, it would have been beneficial for roaming individuals to mix in and mimic, in behavior and appearance, a coral reef fish species unpalatable for a wide number of small-to-medium sized hamlet predators and with simple color patterns. In general, adult butterflyfish in the Caribbean are not a common occurrence in stomach content records (Randall 1967), a testament to their effective defenses, mainly high maneuverability, extremely deep bodies with long, robust spines, particularly in benthivore species, a challenge for their gape-limited predators (Hodge et al 2018). In addition, *C. capistratus* sports less complex color patterns than other Caribbean butterflyfish, an easier starting point for the development of a hamlet mimic.

In conclusion, based on our findings, together with Picq et al (2019) observations, we propose that butter hamlets’ color patterns are unlikely to be the result of direct selection for aggressive mimicry but rather evolved in the context of defensive (Batesian) mimicry, i.e. for predator avoidance. This would configure a ‘social trap’ scenario, as proposed by Robertson (2013) whereby the prey is not the agent of natural selection on aggressive mimicry and resemblance evolves in other contexts, to be later recruited, once evolved, for other functions, in our case aggressive mimicry. Robertson (2013) went so far as to imply that butter hamlet color patterns are not at present under direct selection for aggressive mimicry but simply an unintended by-product of ‘coincidental look-alikes’ with no advantage gained by the inadvertent mimic. Our view is that while the close resemblance between butter hamlet and foureye butterflyfish did not evolve to deceive the visual system of the prey, and therefore not in the context of aggressive mimicry, the resemblance resulting from selection for protective mimicry did eventually start giving butter hamlets an advantage (Puebla et al. 2007) in accessing their prey. While not exerting direct selection on the butter hamlet’s color patterns, this fitness advantage is likely a big contributor to the maintenance of this color morph. Indeed, a defensive and an aggressive role for hamlet mimicry are of course not mutually exclusive and indeed are likely to be both contributing to the maintenance of this behavioral strategy. However, our results suggest that the main prey of butter hamlets, epibenthic masked gobies and mysid shrimp, might have not been the intended signal receiver driving the evolution of mimic coloration, given that both its main prey species are likely devoid of any color vision. On the contrary, predators of hamlets, such as groupers and snappers, while lacking UV vision (Loew and Lythgoe 1978; Losey et al. 2003) that would potentially differentiate model from mimic (Figure 2), are generally at least fully dichromatic when not trichromatic, and likely the receivers of *H. unicolor’* mimic coloration.

Our study shows how the study of sensory systems not only broadens our understanding of animal communication and signaling but has the potential to generate new hypotheses on the origin and maintenance of phenotypic diversity and the evolutionary trajectory of species.

## Supporting information

Suppl. Mat.

