## Supplementary material for "The visual ecology of a color polymorphic reef fish: the role of aggressive mimicry": Suppl. Mat.

Figure S1. Left: Average diffuse attenuation coefficient $\overline{K}_d(z,\lambda)$ (over the 0-7.5m depth interval, above hamlet territories, at Punta Caracol, Bocas del Toro, Panama; *Right*: Spectrum bandwidth (Δλ) of down-, up- and averaged side-welling irradiance, from below surface to close to bottom of reef (7.5m).

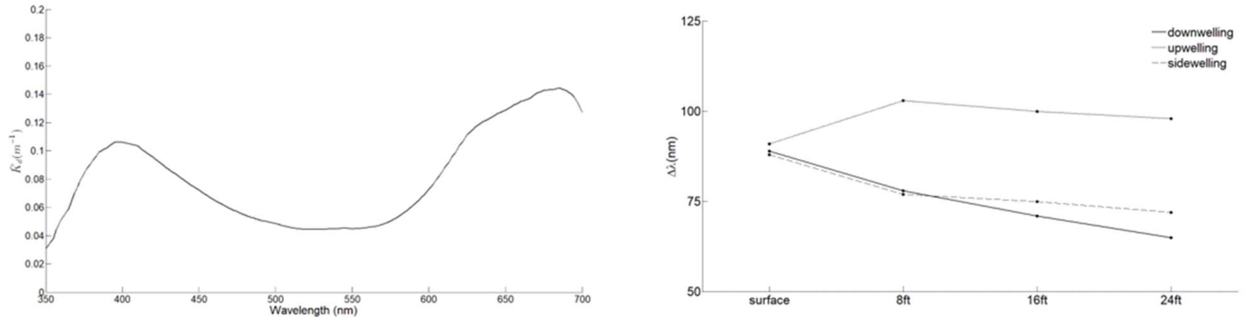

Figure S2. Color distances dS (on the left, in each box) and achromatic distances dL (on the right, in each box) and their bootstrap confidence limits, for each species pair viewed by the prey, a masked goby with a 531/539nm set of cone pigments, at a depth of 5m. A value of dS (or dL) = 1, marks the perceptual threshold, below which a particular patch (#1-7) is likely indistinguishable by the viewer. Figure conventions as in Fig. 4, main text.

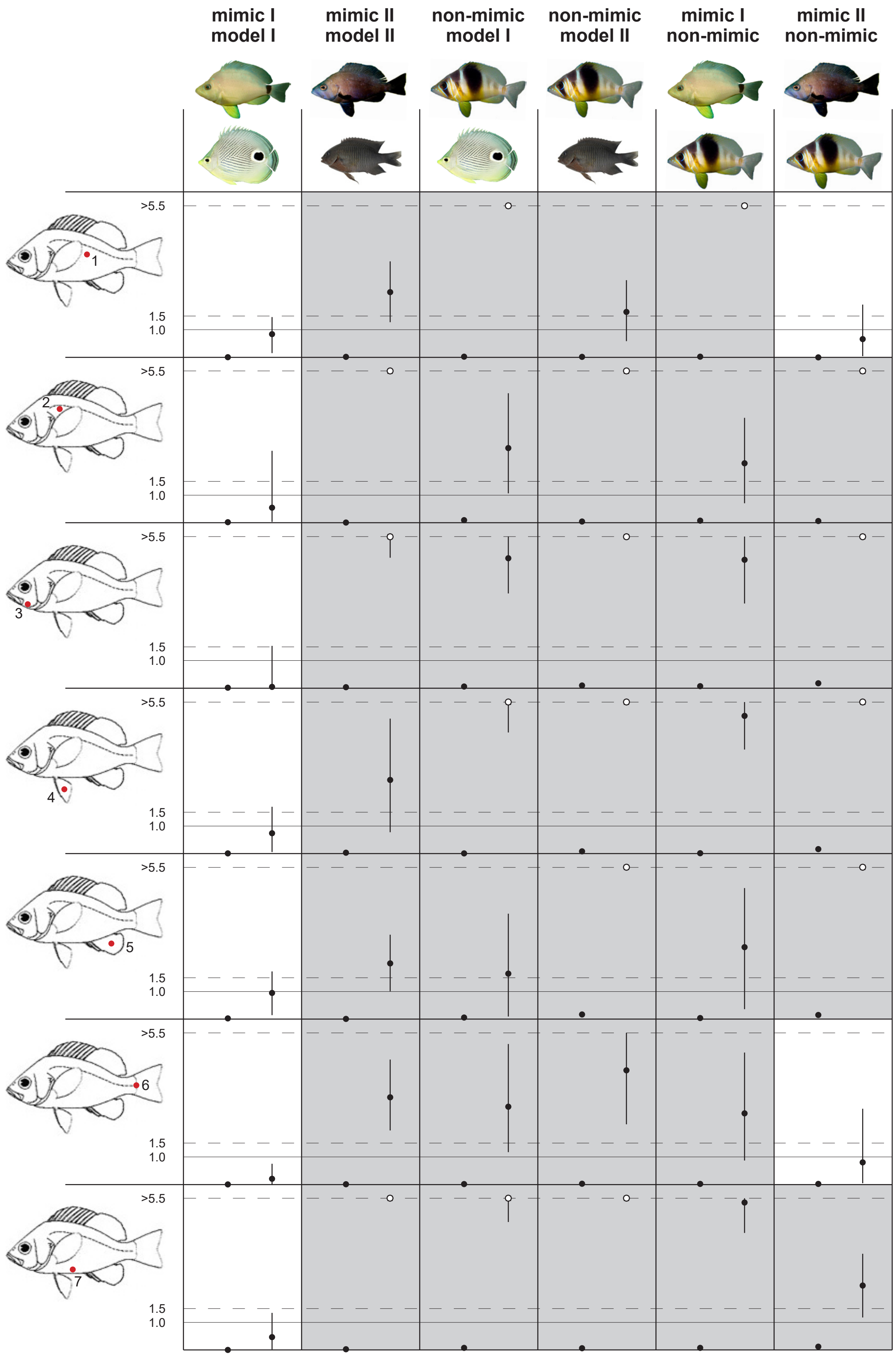

Figure S3. Achromatic contrasts dL across species, viewed by a *Mysidium columbiae* shrimp visual system with a single pigment of $\lambda_{max}$ = 520, at a depth of 5m. Conventions as in Figure 4, main text.

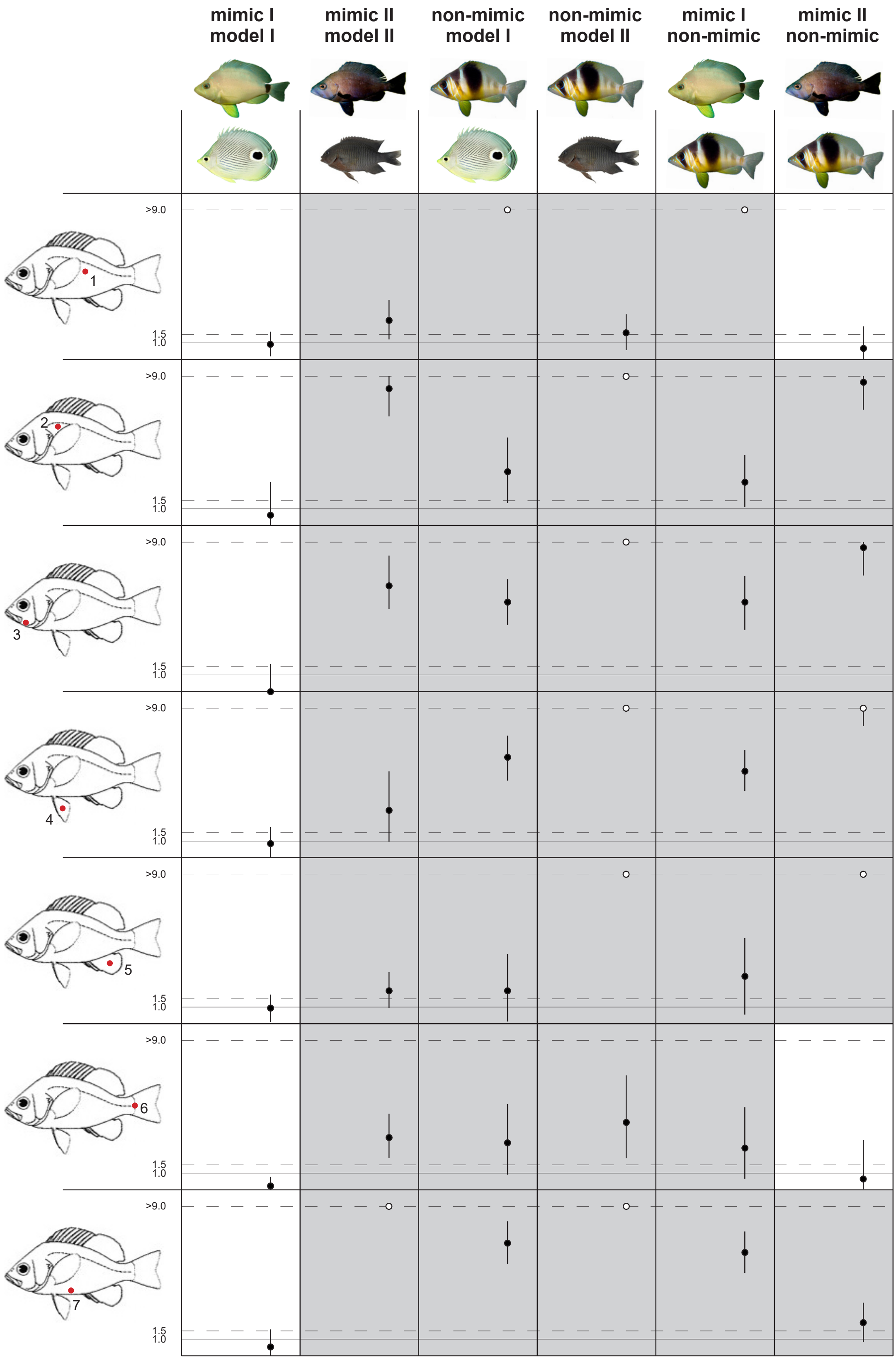

Figure S4. Four-eye butterflyfish (*C. capistratus*), butter hamlet (*H. unicolor*), black hamlet (*H. nigricans*) and barred hamlet (*H. puella*), seen 45° from below (*left*). Detail of ventral appearance (*right*).

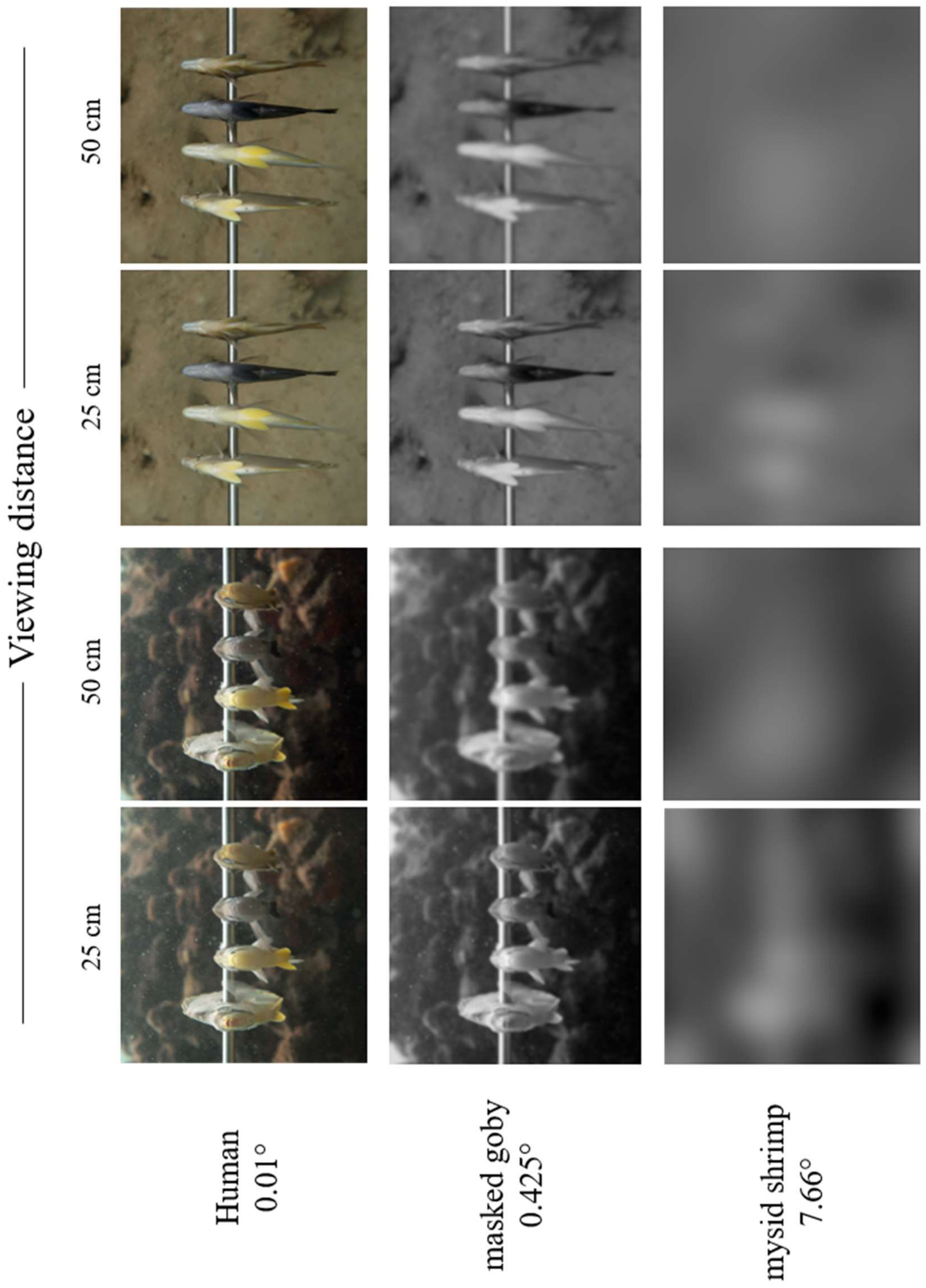

Table S1. Distribution of cone visual pigments in the gobiid species examined to date.

| Species | ROD | SWS | MWS | LWS | habitat | Reference |
|---|---|---|---|---|---|---|
| *Gobiusculus flavescens* | 508 | 453 | 531 | 557 | temperate reef | *Utne-Palm et al 2006* |
| *Pomatoschistus minutus* | 508 | 447 | 527 | 548 | temperate reef | *Jokela-Maatta et al 2009* |
| *Gobius paganellus* | 512 | 465 | 565 | 565 | temperate reef | *Loew and Lythgoe 1978* |
| *Asterropteryx semipunctata* | 498 | — | 531 | 538 | tropical coral reef | *Losey et al 2003* |
| ***Coryphopterus personatus*** | **501** | — | **531** | **539** | **tropical coral reef** | ***this study*** |
